# Flv3A facilitates O_2_ photoreduction and affects H_2_ photoproduction independently of Flv1A in diazotrophic *Anabaena* filaments

**DOI:** 10.1101/2021.12.15.472848

**Authors:** A. Santana-Sánchez, L. Nikkanen, G. Toth, M. Ermakova, S. Kosourov, J. Walter, M. He, E-M. Aro, Y. Allahverdiyeva

## Abstract

The model heterocyst-forming filamentous cyanobacterium, *Anabaena* sp. PCC 7120 (*Anabaena*) represents multicellular organisms capable of simultaneously performing oxygenic photosynthesis in vegetative cells and the O_2_-sensitive N_2_-fixation inside the heterocysts. The flavodiiron proteins (FDPs) have been shown to participate in photoprotection of photosynthesis by driving excess electrons to O_2_ (Mehler-like reaction). Here, we addressed the physiological relevance of the vegetative cell-specific Flv1A and Flv3A on the bioenergetic processes occurring in diazotrophic *Anabaena* under variable CO_2_. We demonstrate that both Flv1A and Flv3A are required for proper induction of the Mehler-like reaction upon a sudden increase in light intensity, which is likely important for the activation of carbon-concentrating mechanisms (CCM) and CO_2_ fixation. Under ambient CO_2_ diazotrophic conditions, Flv3A is capable of mediating moderate O_2_ photoreduction, independently of Flv1A, but in coordination with Flv2 and Flv4. Strikingly, the lack of Flv3A resulted in strong downregulation of the heterocyst-specific uptake hydrogenase, which led to enhanced H_2_ photoproduction under both oxic and micro-oxic conditions. These results reveal a novel regulatory network between the Mehler-like reaction and the diazotrophic metabolism, which is of great interest for future biotechnological applications.

## Introduction

Filamentous heterocyst-forming cyanobacteria such as *Anabaena* sp. PCC 7120 (hereafter *Anabaena*) represent a unique group of prokaryotes capable of simultaneously performing two conflicting metabolic processes: (i) O_2_-producing photosynthesis in vegetative cells; and (ii) O_2_-sensitive N_2_ fixation in heterocysts. This ability has evolved through cellular differentiation under nitrogen limiting growth conditions when some vegetative cells from the filament transform into specialized heterocyst cells that provide a microaerobic environment suitable for N_2_ fixation. H_2_ gas is naturally produced as an obligatory by-product of the N_2_-fixation process carried out by nitrogenase, which is highly sensitive to O_2_. The natural yield of H_2_ gas production inside heterocysts is limited. This is due to rapid H_2_ recycling, mainly by an uptake hydrogenase enzyme, which further returns electrons for the N_2_-fixing metabolism (Tsygankov et al., 2007; Bothe et al., 2010).

In oxygenic photosynthesis, light drives the photosynthetic linear electron transport from water to NADPH, using Photosystem (PS) II, Cytochrome (Cyt) *b*_*6*_*f* and PSI complexes embedded in the thylakoid membrane. These electron transport reactions are coupled to ATP synthesis *via* the generation of trans-thylakoid proton motive force (*pmf)*. The obtained NADPH and ATP are then used as reducing power for CO_2_ fixation and cell metabolism. Environmental fluctuations in light and nutrient supply might result in the over-reduction of the photosynthetic machinery. Alleviation of excess electrons by the class-C Flavodiiron proteins (hereafter FDP) has been described in all oxygenic photosynthetic organisms, apart from angiosperms, red and brown algae (Helman et al., 2003; Zhang et al., 2009; Jokel et al., 2018; Gerotto et al., 2016; Chaux et al., 2017; Ilik et al., 2017; Alboresi et al., 2019; Shimakawa et al., 2021). This group of proteins act as strong electron outlets downstream of PSI by catalyzing the photoreduction of O_2_ into H_2_O (named the Mehler-like reaction) (Allahverdiyeva et al., 2011; 2013; 2015; Santana-Sánchez et al., 2019).

Six genes encoding FDPs have been reported in *Anabaena* (Ow et al., 2008; Zhang et al., 2009; Ermakova et al., 2013; Allahverdiyeva et al., 2015). Phylogenetic assessment has shown that four of these genes (*flv1A, flv3A, flv2*, and *flv4*) are highly similar to their homologs in *Synechocystis*. sp. PCC 6803 (hereafter *Synechocystis*), SynFlv1-SynFlv4. Recently, we demonstrated that SynFlv1 and SynFlv3 proteins function in coordination with, but distinctly from SynFlv2 and SynFlv4 (Santana-Sánchez et al., 2019). While the SynFlv1/Flv3 hetero-oligomer is mainly responsible for the initial fast and transient O_2_ photoreduction during a sudden increase in light intensity, SynFlv2/Flv4 catalyzes steady O_2_ photoreduction under illumination at air-level CO_2_ (LC). Importantly, the single deletion of any FDP strongly diminishes the O_2_-photoreduction, indicating that O_2_ photoreduction is mainly catalyzed by the hetero-oligomeric forms working in an interdependent manner (Santana-Sánchez et al., 2019; Nikkanen et al., 2020).

The two additional *Anabaena* FDP proteins, AnaFlv1B and AnaFlv3B, are exclusively localized in the heterocysts (Ermakova et al., 2013). The AnaFlv3B protein was shown to mediate the photoreduction of O_2_ independently of AnaFlv1B, likely as a homo-oligomer, playing an important role in maintaining micro-oxic conditions inside heterocysts under illumination, which is crucial for N_2_ fixation and H_2_ production (Ermakova et al., 2014). However, research on the role of heterocyst-specific AnaFlv1B and vegetative cell-specific FDPs in diazotrophic cyanobacteria is still scarce.

Here, we addressed the physiological relevance of the AnaFlv1A and AnaFlv3A isoforms on the bioenergetic processes occurring in vegetative cells and heterocysts of diazotrophic *Anabaena*. AnaFlv1A and AnaFlv3A were shown to have a crucial photoprotective role under fluctuating light intensities (FL), regardless of nitrogen or CO_2_ availability, suggesting functional analogy with homologs in *Synechocystis*. Importantly however, our results also provided evidence for distinct functional roles of AnaFlv3A and AnaFlv1A. We showed that by cooperating with AnaFlv2 and/or AnaFlv4, AnaFlv3A can function independently of AnaFlv1A in O_2_ photoreduction in low CO_2_ conditions. AnaFlv3A was also indirectly linked with the H_2_ metabolism occurring inside heterocyst cells. Our work highlights the complex regulatory network between oxygenic photosynthesis, nitrogen fixation and hydrogen photoproduction.

## Materials and Methods

### Strains and culture conditions

*Anabaena* sp. PCC 7120 strain was used as the wild-type (WT) in this study. The Δ*flv1A* and Δ*flv3A* mutants (Allahverdiyeva et al., 2013) and the Δ*hupL* mutant (Masukawa et al., 2002) were previously reported. For construction of the double mutant Δ*flv1A/flv3A*, the BamHI-XbaI region of the mutated *flv1A* construct was replaced with the spectinomycin/streptomycin resistance cassette. The generated plasmid was transferred into Δ*flv3A* and sucrose, neomycin, and spectinomycin was used for selection. Segregation of the mutant was verified by PCR. Culture stocks of Δ*flv1A* and Δ*flv3A* mutants were maintained in BG-11 medium supplemented with 40 μg mL^-1^ neomycin, while the Δ*hupL* mutant was supplemented with 20 μg mL^-1^ spectinomycin.

Pre-cultures were grown in Z8x medium (lacking combined nitrogen, pH 7.0-7.3, Kotai, 1972) at 30 °C and under constant white light of 75 μmol photons m^−2^ s^−1^ without antibiotics. For this, the filaments were inoculated at OD_750_= 0.1 in 200 mL Z8x medium (in 500 mL flasks) and were continuously bubbled with air (0.04% CO_2_, LC) or with air supplemented with 1% CO_2_ (HC) if not specifically mentioned. Pre-cultures were harvested at the logarithmic growth phase, inoculated at OD_750_= 0.1 in fresh Z8x medium and experimental cultures were grown under similar pre-experimental conditions (75 μmol photons m^−2^ s^−1^ illumination and bubbling with air or 1% CO_2_ supplemented). Experimental cultures were harvested after 4 days of growth and experiments were conducted in 3-5 independent biological replicates.

### Determination of heterocyst frequency

Alcian blue was used to stain the polysaccharide layer of the heterocyst envelope. Cell suspensions were mixed (1:8) with a solution of 0.5% Alcian Blue stain in 50% ethanol-water. Stained samples were visualized using a Wetzlar light microscope (Leitz) and x400 magnification micrographs were taken. Around 1000-2000 cells were counted per sample, and the heterocyst frequency was determined as a percentage of total cells counted.

### MIMS measurements

*In vivo* measurements of ^16^O_2_ (m/z = 32), ^18^O_2_ (m/z = 36), CO_2_ (m/z = 44) and H_2_ (m/z = 2) fluxes were monitored using a membrane inlet mass spectrometry (MIMS) as described previously (Mustila et al., 2016). Harvested filaments were resuspended with fresh Z8x medium, adjusted to Chl *a* 10 μg mL^−1^ and acclimated for 1 hr to the growth conditions. For LC samples, the concentration of dissolved total inorganic carbon was saturated with 1.5 mM NaHCO_3_ before the measurement.

To measure Deuterium uptake, the filaments were flushed with Ar inside gas-tight vials for 15 min, then 1.2 mL pure D_2_ (2 % in headspace) was injected into each vial. Changes of D_2_ in the gas phase were measured at 2 h and 24 h after D_2_ addition. 250 μL gas sample from the headspace of the vials was injected into the MIMS chamber. The calibration of D_2_ concentration was performed by injecting known concentrations of D_2_ into the media.

### Fluorescence analysis

A pulse amplitude modulated fluorometer Dual-PAM-100 (Walz) was used to monitor Chl *a* fluorescence and P700 absorbance. Harvested filaments were resuspended in fresh Z8x medium to the Chl *a* concentration of 15 μg mL^-1^ and then kept for about 1 hr under the growth conditions. Before the measurements, samples were dark-adapted for 10 min. The measurement started with a saturating pulse in darkness to determine F_m_^D^. Then, the samples were illuminated with red actinic light at a photon flux density of 50 μmol photons m^-2^ s^-1^ for 380 s whilst saturating pulses (SP, 5000 μmol photons m^−2^ s^−1^, 400 ms) were given every minute (SP1-SP9). Photosynthetic parameters were determined as described previously (Huokko et al 2017).

### Determination of PSI primary donor (P700) and Ferredoxin (Fd) redox changes from near-infrared absorbance

The absorbance differences at 780–820 nm, 820–870 nm, 840–965 nm and 870–965 nm were measured with the Dual KLAS/NIR spectrophotometer (Walz). Experimental cultures, as well as cultures used for determination of the model spectra (more info in Supplementary Mathods), were grown at 50 μmol photons m^-2^ s^-1^ and under air-level CO_2_ (LC) in Z8x medium for 4 days, then adjusted to *Chl* a concentration of 20 μg mL^-1^ by re-inoculating pelleted cells in fresh medium. Cells were dark-adapted for 10 min, after which absorbance differences of the four wavelength pairs were measured during 5 s actinic illumination at 500 μmol photons m^-2^ s^-1^ and subsequent dark. The maximal levels of P700 oxidation and Fd reduction were determined for each sample by utilizing the NIRMAX script (Klughammer and Schreiber 2016), and the obtained experimental deconvoluted traces were then normalized to the maximal values. The Dual-KLAS/NIR measurement of *Synechocystis* Δ*flv1* cells was performed as described previously (Nikkanen et al., 2020).

### H_2_ measurement by Clark-type electrode

H_2_ concentration was monitored under anaerobic conditions using a Clark-type Pt-Ag/AgCl electrode chamber (DW1/AD, Hansatech) connected to a homemade polarographic box. Experimental cultures were harvested, resuspended in fresh Z8x medium and adjusted to the Chl *a* concentration of about 3-4 μg mL^-1^. The resulting suspensions (∼30 mL) were transferred into 75 mL glass vials, sealed and sparged with either nitrogen (N_2_) or argon (Ar) for 30 min in the dark to achieve anaerobic conditions. Then, cultures were incubated under the corresponding atmosphere for another 2 h in the dark at 25 °C. 4 mL of dark-adapted suspension were transferred into the chamber with an anaerobic gas-tight syringe and H_2_ concentration was monitored during 6 min illumination with actinic light of 800 μmol photons m^−2^ s^−1^ after which the light was switched off. The H_2_ production rates were calculated using linear regression.

### Nitrogenase activity essay

Acetylene reduction assay was used to determine nitrogenase activity as described previously (Leino et al. 2014). 5 mL of experimental samples were transferred into 23 mL vials, flushed with argon for 20 min and supplemented with 10% acetylene in the headspace. Vials were kept for 20 hr under 50 μmol photons m^−2^ s^−1^ at 30 °C with gentle agitation (120 rpm). After this, 20 μL of gas sample was withdrawn from the headspace of the vial and analysed for ethylene content using a gas chromatograph equipped with Carboxen(r)-1010 PLOT Capillary Column and FID detector. The enzyme activity was calculated from the peak area and normalised to the total protein content.

### Chl *a* and total sugar determination

Chl *a* was extracted from cells in 90% methanol and the concentration was determined by measuring absorbance at 665 nm and multiplying it with the extinction coefficient factor 12.7 (Meeks & Castenholz, 1971). For total sugar determination, 1 mL of experimental samples were collected, washed and diluted to 1:1 with MQ-water before the sugar measurement. Total sugar content was obtained using the colourimetric method described earlier (DuBois et al., 1956).

### Protein extraction and immunoblotting

Total protein extracts were isolated as described previously (Zhang et al., 2009). Electrophoresis and immunoblotting were performed according to an earlier report (Mustila et al., 2016). Protein-specific antibodies raised against Flv3A (Agrisera), PsaB (AS10 695, Agrisera), NdhK (Agrisera), and HupL (kindly provided by P. Tamagnini) were used in this study.

### RNA isolation and RT-qPCR analysis

Isolation of total RNA, reverse transcription and qPCR analysis was performed as described earlier (Ermakova et al., 2013). *rnpB gene* was used as a reference for normalization. The primer pairs used in this study are listed in Table S1.

## Results

### Phenotypic characterization of *Anabaena* mutants deficient in Flv1A and Flv3A

To investigate the function of the vegetative cell-specific Flv1A and Flv3A proteins in diazotrophic *Anabaena* filaments, we used Δ*flv1A* and Δ*flv3A* deletion mutants (Fig. S1 and Allahverdiyeva et al., 2013). Likewise, the SynFlv1 and SynFlv3, the AnaFlv1A (encoded by *all3891*) and AnaFlv3A (encoded by *all3895*) proteins are indispensable for diazotrophic and non-diazotrophic growth of *Anabaena* filaments under severe fluctuating light intensities at both air level (low CO_2_, LC) and 1-3 % CO_2_ (high CO_2_, HC) (Fig. S2a, Allahverdiyeva et al., 2013).

Under constant light (at a photon flux density of 50 μmol photons m^−2^ s^−1^), there were no significant differences in the growth of these mutants compared to the WT, as measured by OD_750_ (Fig. S2b) or concentration of chlorophyll *a* (Chl *a*). Total protein and sugar content of the WT and Δ*flv1A* and Δ*flv3A* filaments were also similar (Table 1). Light microscopic images of *Anabaena* filaments indicated that both Δ*flv1A* and Δ*flv3A* mutants and WT had a similar ratio of vegetative cells to heterocysts (Table 1) and no visible changes were observed in heterocyst morphology.

**Table 1.** Growth characteristics and photosynthetic parameters of the WT, Δ*flv1A*, and Δ*flv3A* filaments. Experimental cultures were grown under diazotrophic LC conditions for 4 days. The maximum quantum yield of PSII (Fv/Fm), minimal level of fluorescence (F_o_), maximal fluorescence in the dark (F_m_^D^), maximal fluorescence (F_m_^’^), quenching due to state transition (qT). Values are means ± SD, n = 3-5 biological replicates. Asterisks indicate statistically significant differences compared to the WT (t-test, *P* < 0.05).

| Parameters | WT | $\Delta flv1A$ | $\Delta flv3A$ |
| --- | --- | --- | --- |
| OD<sub>750</sub> | 1.46±0.18 | 1.45±0.13 | 1.43±0.18 |
| Chl *a*, µg mL<sup>-1</sup> | 7.01±0.80 | 6.66±0.76 | 6.97±0.96 |
| Total protein, µg mL<sup>-1</sup> | 263.6±31.7 | 265.3±10.1 | 254±14.1 |
| Total sugars, µg mL<sup>-1</sup> | 53.8±12.7 | 45.4±4.9 | 43.9±5.8 |
| Heterocyst frequency (%) | 12.7±1.1 | 12.1±0.9 | 10.7±2.1 |
| Nitrogenase activity (C<sub>2</sub>H<sub>2</sub> reduction),<br>µmol mg Chl<sub>a</sub><sup>-1</sup>h<sup>-1</sup> | 21.3±4.0 | 16.7±4.6 | 15.9±2.7* |
| F<sub>v</sub>/F<sub>m</sub> (with 10 µM DCMU) | 0.45±0.05 | 0.5±0.01 | 0.49±0.01 |
| F<sub>o</sub> | 0.91±0.01 | 0.86±0.00* | 0.84±0.00* |
| F<sub>m</sub><sup>D</sup> | 1.39±0.02 | 1.25±0.02* | 1.26±0.01* |
| F<sub>m</sub>' (SP 1) | 1.46±0.05 | 1.42±0.04 | 1.49±0.04 |
| qT | 0.05±0.02 | 0.14±0.01* | 0.18±0.01* |

### Fluorescence and P700 oxidoreduction properties of Δ*flv1A* and Δ*flv3A*

Diazotrophic *Anabaena* WT, Δ*flv1A*, and Δ*flv3A* filaments, grown under LC and HC, were next subjected to fluorescence analyses under growth light intensity to disclose the impact of Flv1A and Flv3A, common or specific, on photosynthetic electron transport. The dark-adapted WT filaments showed a relatively low maximal fluorescence in the dark (F_m_^D^) (state 2). Upon exposure to actinic light intensity, maximal fluorescence (F_m_’) slightly increased, indicating a transition of filaments to state 1 (Fig. 1a). The state 2-to-state 1 transition observed upon illumination was less pronounced in HC-grown WT filaments (Fig. 1b). The effective yield of PSII [Y(II)], calculated for each saturating pulse (SP), remained stable (0.34±0.02 - 0.30±0.01) during the illumination of WT filaments grown both under LC (Fig. 1c) and HC (Fig. 1d) conditions.

**Figure 1.**
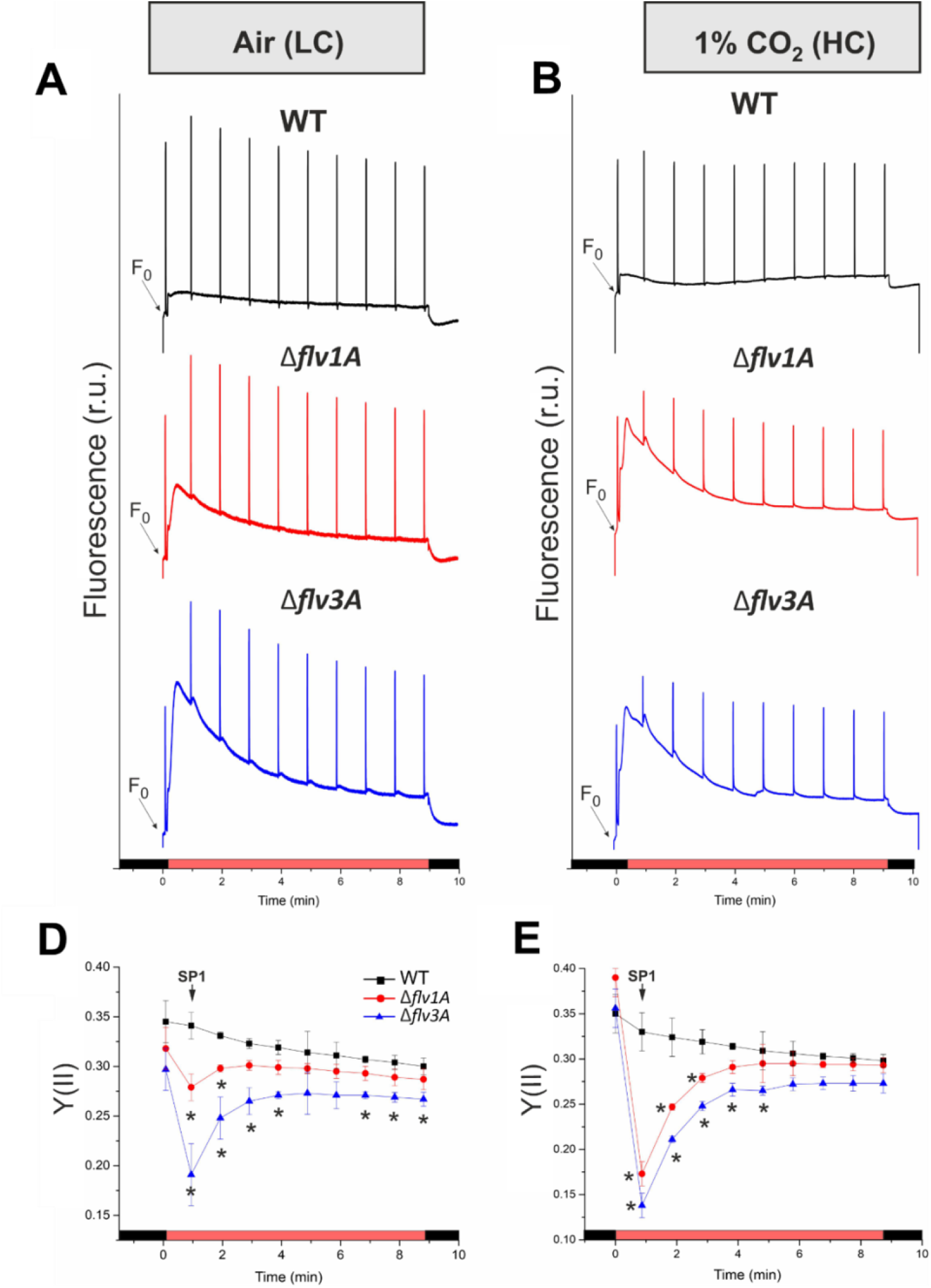
Fluorescence analysis of the diazotrophic *Anabaena* WT, *Δflv1A*, and *Δflv3A*. (A, C) Filaments were cultivated under air (LC) or (B, D) under air supplemented with 1% CO_2_. (HC). Representative traces of 3 biological replicates are shown (A, B). Cells were dark acclimated for 10 min before illumination with 50 μmcl photons m^-2^ s^-1^ of actinic light- The effective yield of PSII [Y(ll)] was calculated by applying saturating pulses during induction curve measurements (C, D). Values are means ± SD, n = 3 biological replicates. Asterisks indicate statistically significant differences compared to the WT (t-test, *P*<0.05). r.u., relative units.

Both the Δ*flv1A* and Δ*flv3A* mutants showed significantly lower F_m_^D^ than WT in LC (Fig. 1 and Table 1), implying a more pronounced state 2 in the dark. Accordingly, a stronger state 2-to-state 1 transition (qT in Table 1) was observed during illumination in comparison to WT, similarly to the phenotype previously described in the *Synechocystis Δflv3* mutant (Elanskaya et al., 2021). Notably, during the dark-to-light transition, the fluorescence kinetics were differently affected in the two mutants grown under LC conditions. Illumination of Δ*flv3A* filaments resulted in a rapid increase of the fluorescence level which was gradually quenched but remained at a higher steady-state level (F_s_) compared to WT and the Δ*flv1A* mutant. The Δ*flv1A* mutant showed only a moderate increase and then a gradual decay of fluorescence, reaching the WT F_s_ level after 4 min of illumination. Differently from LC grown filaments, the Δ*flv1A* and Δ*flv3A* mutants grown under HC conditions revealed a similar fluorescence increase during the dark-to-light transition, which gradually decayed and reached the WT levels by the end of the illumination period (Fig. 1b).

The effective yield of PSII, Y(II) echoed high fluorescence levels and small variable fluorescence (F_v_’) upon illumination by SP1, showing a strong drop both in LC-(84% and 62% that of WT in Δ*flv1A* and Δ*flv3A* mutants, respectively) and HC-grown filaments (53% and 39% that of WT in Δ*flv1A* and Δ*flv3A* mutants, respectively) (Fig. 1c,d). After that, the Y(II) values gradually recovered over the course of illumination, though Δ*flv3A* did not reach the WT levels (Fig. 1c). Notably, the maximum quantum yield of PSII, F_v_/F_m_, did not differ significantly between the mutants and the WT (Table 1).

Examination of the transient post-illumination increase of fluorescence level (F_0_ rise), which reflects the NDH-1 mediated reduction of the PQ pool in darkness (Mi et al., 1995), demonstrated a notably higher F0 rise in both Δ*flv1A* and Δ*flv3A* mutants grown under LC and HC conditions (Fig. S3). In line with the lower FmD (Table 1), this finding suggests elevated electron flux into the PQ pool in the Δ*flv1A* and Δ*flv3A* mutants, presumably mediated by NDH-1, in comparison to WT. Considering that the abundance of NdhK, the core subunit of the NDH-1 complex, was similar between all genotypes (Fig. S4d) the difference in F_0_ rise is likely caused by higher availability of reduced ferredoxin (Fd), the likely electron donor to both FDPs and NDH-1 (Nikkanen et al., 2021), or by post-translational regulatory factors.

At the onset of high irradiance both the Δ*flv1* (Fig. S5) and Δ*flv3* mutants of *Synechocystis* are unable to rapidly re-oxidize Fd, causing accumulation of electrons at P700 (Nikkanen et al., 2020; Theune et al., 2021). To determine whether this occurs in the *Anabaena* Δ*flv1A* and Δ*flv3A* mutants, we determined the high light-induced fast redox changes of Fd and P700 from near-infrared absorbance differences using the Dual KLAS/NIR spectrophotometer. The results indicated that similarly to *Synechocystis* Δ*flv3* (Nikkanen et al., 2020) and Δ*flv1* mutants (Fig. S5), both *Anabaena* mutants also suffered from delayed re-oxidation of Fd and P700 upon illumination, and showed slower post-illumination re-oxidation of Fd (Fig. 2). Unlike in the *Synechocystis* mutants, there was a clear difference between the Δ*flv1A* and Δ*flv3A* mutants of *Anabaena*, with the Δ*flv3A* strain displaying more severe delay in re-oxidation of Fd and P700 than Δ*flv1A*.

**Figure 2.**
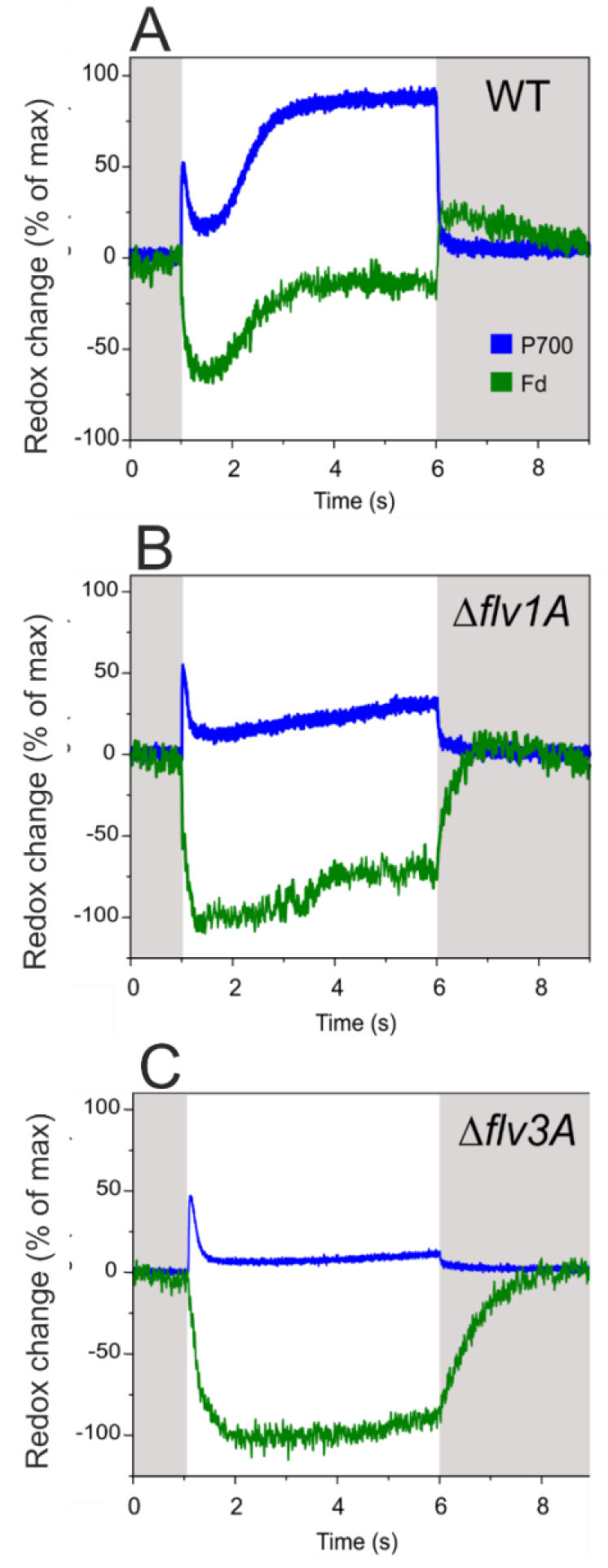
Redox changes of P700 and Fd upon dark-light-dark transitions in the diazotrophic *Anabaena* WT, Δ*flvIA*, and Δ*flv3A* filaments. The cells were grown under 50 μmol photons m^-2^ s^-1^ and air-level CO_2_ (LC) for 4 days, harvested and adjusted to Chl a concentration of 20 μg mL^-1^. Cells were dark-adapted for 10 min, after which absorbance differences of four near-infrared wavelength pairs were measured with the Dual KLAS/NIR spectrophotometer during 5 s actinic illumination at 503 μmol photons m^-2^ s^-1^ and subsequent darkness. P700 and Fd redox changes were then deconvolved from the absorbance differences using specifically determined differential model plots (model spectra) for *Anabaena* (see Materials and methods). Maximal levels of Fd reduction and P700 oxidation in each sample were used to normalize the traces. Representative traces of 3 biological replicates are shown.

### Real-time gas exchanges in *Anabaena* FDP mutants

To clarify the specific impacts of *flv1A* and *flv3A* deletions on real-time gas fluxes in the diazotrophic filaments of *Anabaena*, we used membrane inlet mass spectrometry (MIMS) analysis (Fig. 3). The MIMS technique combined with the use of ^18^O_2_ isotopologue allows distinguishing between light-induced O_2_ reduction (uptake) and photosynthetic O_2_ production. The net O_2_ evolution rate was calculated as the difference between the rates of gross O_2_ evolution and O_2_ uptake in the light. For all MIMS measurements, gas exchange was monitored for 4 min in dark followed by 5 min of high irradiance (500 μmol photons m^-2^ s^-1^) and for an additional 3 min in the dark.

**Figure 3.**
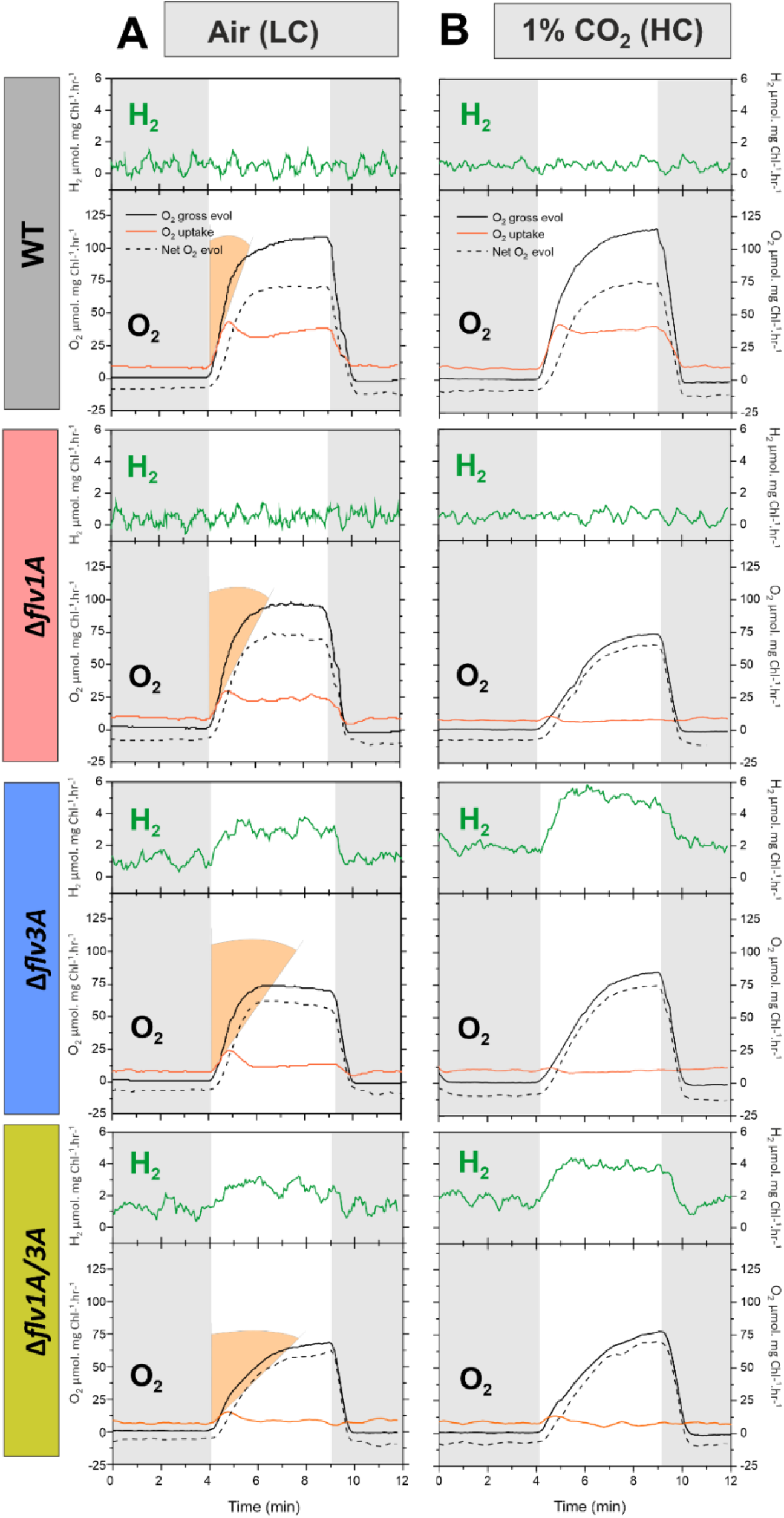
O_2_ and H_2_ exchange rates of the diazotrophic *Anabaena* WT, Δ*flv1A* and Δ*flv3A* filaments. The filaments were cultivated for 4 days under air (LC) (A) and high carbon (1 % CO_2_ in the air, HC) (B), after which the filaments were harvested and the Chl *a* concentration adjusted to 10 μg mL^-1^. Gas exchange rates were calculated in darkness (grey areas of the graphs) and under illumination with actinic white light at 500 μmol photons m^-2^ s^-1^. For LC measurements, samples were supplemented with 1.5 mM NaHCO_3_. Orange shading indicates the differences in the initial slope of the O_2_ photoreduction rates. The plots are representative of three independent biological replicates.

Illumination of WT filaments grown under LC demonstrated a rapid increase in the rate of O_2_ uptake from 10.6±2.7 μmol O_2_ mg Chl *a*^-1^ h^-1^ in darkness to 34.6±0.2 μmol O_2_ mg Chl *a*^-1^ h^-1^ in light. This fast induction phase was followed by a decay that stabilized at 28.4±1.4 μmol O_2_ mg Chl *a*^-1^ h^-1^ after 3 min (Fig. 3a, Table S2). This pattern resembles, to some extent, previously described triphasic kinetics of O_2_ photoreduction in *Synechocystis* grown under LC conditions (Santana-Sánchez et al., 2019), and in HC-grown *Chlamydomonas reinhardtii* cells illuminated with high light intensity (Saroussi et al., 2019). Both mutants showed slightly lower O_2_ uptake rates in darkness than the WT (Table S2) but the rate of O_2_ consumption under illumination was affected to different extents in Δ*flv1A* and Δ*flv3A* filaments. The Δ*flv3A* mutant exhibited strong impairment of light-induced O_2_ uptake, showing a maximal rate of 15.6±0.1 μmol O_2_ mg Chl *a*^-1^ h^-1^ (52 % lower than the WT) at the onset of light, which declined to a residual rate of 4.3±0.2 μmol O_2_ mg Chl *a*^-1^ h^-1^ by the end of illumination. In contrast to the *Synechocystis* Δ*flv1* mutant (Fig. S7) where O_2_ reduction is almost fully eliminated, the *Anabaena* Δ*flv1A* filaments showed an intermediate phenotype whereby a maximum light-induced O_2_ reduction rate of 22.2±2.8 μmol O_2_ mg Chl *a*^-1^ h^-1^ (34% lower than WT) was observed, which declined to 15.5±1.2 μmol O_2_ mg Chl *a*^-1^ h^-1^ (Fig. 3a). Moreover, both Δ*flv1A* and Δ*flv3A* mutants showed slower activation of O_2_ photoreduction, with a more pronounced lag-phase in Δ*flv3A* (Fig. 3a, orange shading). These results suggested that both AnaFlv1A and AnaFlv3A contribute to the Mehler-like reaction, but to a differing extent and presumably in different homo/hetero-oligomeric arrangements.

To clarify whether the homo-oligomers of AnaFlv1A in Δ*flv3A* and conversely, the homo-oligomers of AnaFlv3A in Δ*flv1A* mutants contribute to the observed O_2_ photoreduction rates (Fig. 3a), we constructed a double mutant Δ*flv1A*/Δ*flv3A* (Fig. S6). MIMS analysis revealed that concomitant inactivation of both *flv1A* and *flv3A* strongly inhibited the O_2_ photoreductionin *Anabaena* filaments cultivated under LC conditions (Fig. 3a) suggesting either contribution of AnaFlv1 and AnaFlv3 homo-oligomers to O_2_ photoreduction in the single mutants or involvement of AnaFlv2 and/or AnaFlv4 proteins in this process. Previous studies with *Synechocystis* cells (Zhang et al., 2009; Wang et al., 2004; Eisenhut et al., 2012; Santana-Sánchez et al., 2019) and non-diazotrophic *Anabaena* WT filaments (Ermakova et al., 2013) demonstrated high transcript abundance of *flv2* and *flv4* at LC. Therefore, we next investigated the abundance of *flv2* and *flv4* transcripts in diazotrophic *Anabaena* filaments grown under LC and HC using RT-qPCR. The Δ*flv1A* and Δ*flv3A* mutants grown under LC demonstrated significantly higher *flv2* and *flv4* transcript levels compared to the WT (Fig. S4a). Under HC, transcript abundances of *flv2* and *flv4* did not differ between the mutants and the WT but were drastically lower in all genotypes compared to LC conditions (Fig. S4b). This prompted us to examine the possible contribution of AnaFlv2 and AnaFlv4 proteins to the Mehler-like reaction by comparing the O_2_ photoreduction rates in Δ*flv1A* and Δ*flv3A* mutants grown under LC (Fig. 3a) *vs* HC conditions (Fig. 3b), where the expression of *flv2* and *flv4* were found to be induced and repressed, respectively.

While the O_2_ photoreduction in WT filaments grown under HC was comparable to that under LC conditions (Fig. 3a), the inactivation of *flv1A* and/or *flv3A* fully eliminated light-induced O_2_ reduction in the filaments grown under HC (Fig. 3b). This result suggests that the highly expressed *flv2* and *flv4* likely contribute to O_2_ photoreduction in diazotrophic Δ*flv1A* and Δ*flv3A* filaments grown under LC conditions. Nevertheless, further elucidation is needed to verify the functioning of the AnaFlv2/Flv4 hetero-oligomer or different FDP oligomer compositions in O_2_ photoreduction.

It is important to note that under LC conditions, while gross O_2_ evolution and net photosynthetic O_2_ production rates of the Δ*flv1A* mutant were comparable to those of the WT, the Δ*flv3A* mutant demonstrated lower gross and net O_2_ evolution rates (Fig. 3a, Table S2). Strikingly, the initial peak in CO_2_ uptake rates associated with the CCM activation (Liran et al., 2018) as well as the steady-state of CO_2_ fixation of both deletion mutants were significantly diminished compared to the WT, and the Δ*flv3A* strain showed pronounced impairment than Δ*flv1A* (Table S2, Fig. S8b). Under HC conditions, both mutants had lower gross O_2_ evolution (65.9±7.3 and 76.8±6.5 μmol O_2_ mg Chl *a*^-1^ h^-1^, respectively) relative to the WT (122.2±14.8 μmol O_2_ mg Chl *a*^-1^ h^-1^) and a delay in the induction of O_2_ evolution upon illumination (Fig. 3b). Accordingly, the activation of CO_2_ fixation under HC conditions was slower and decreased in both mutants compared to WT (Fig. S8c). In the Δ*flv1A*/Δ*flv3A* double mutant a delay in gross O_2_ evolution was observed under LC that was more severe than in Δ*flv3A* cells (Fig. 3A), while under HC all three mutant strains were similarly impaired (Fig. 3b). This suggests that not only AnaFlv3A, but also AnaFlv1A may be performing some AnaFlv2-4-dependent but AnaFlv3A-independent function in LC that affects photosynthetic electron transport.

### Consequences of *flv1A* or *flv3A* deletion on diazotrophic metabolism

Based on the results above it is clear that AnaFlv1A and AnaFlv3A impact the photosynthetic apparatus to different extents in LC-grown diazotrophic *Anabaena*. In comparison to the inactivation of AnaFlv1A, the deletion of AnaFlv3A resulted in a stronger reduction of the PQ pool, leading to a consistent decrease of PSII effective yield (Fig. 1c) and, eventually, lower net O_2_ evolution rates over the illumination period (Fig. 3a). On the other hand, previous studies with different diazotrophic *Anabaena* species have demonstrated that the disruption of PSII activity in vegetative cells has implications for N_2_ and H_2_ metabolism inside heterocysts, thus, modulating the diazotrophic metabolism of filaments (Khetkorn et al., 2012; Chen et al., 2014). We, therefore, examined whether the absence of AnaFlv1A or AnaFlv3A from vegetative cells has a long-distance impact on the heterocyst metabolism. To this end, we analyzed the nitrogenase activity and H_2_ fluxes of diazotrophic *Anabaena* WT, Δ*flv1A* and Δ*flv3A* mutants.

As demonstrated in Table 1, both Δ*flv1A* and Δ*flv3A* mutants showed somewhat lower nitrogenase activity in comparison to WT filaments, yet only the nitrogenase activity of the Δ*flv3A* mutant was significantly lower compared to WT. Real-time gas exchange monitored by MIMS (Fig. 3) revealed no changes in the H_2_ gas concentration in WT and Δ*flv1A* during the dark-light transition. In contrast, the Δ*flv3A* mutant showed an increase in H_2_ level in the dark and a clear light-induced H_2_ gas production (1.7±0.4 μmol mg Chl *a*^-1^ h^-1^) (Fig. 3A). This result was confirmed by a second independent Δ*flv3A* mutant strain showing similar light-induced H_2_ production (Δ*flv3A*_C2 in Fig. S8a). Interestingly, the Δ*flv3A* mutant cultivated under HC conditions demonstrated an even higher H_2_ photoproduction rate (2.8±0.8 μmol mg Chl *a*^-1^ h^-1^, Fig. 3b). Although Δ*flv3A* filaments showed real-time H_2_ production under oxic conditions, the rate of H_2_ production remained low.

Next, we monitored H_2_ in anoxic cultures using a Clark-type electrode. Under the N_2_ atmosphere, the Δ*flv3A* mutant demonstrated a significantly higher yield of H_2_ photoproduction accompanied by a three times higher specific H_2_ production rate compared to the WT (Fig. 4a, Fig. S9a). To confirm that the observed H_2_ production is nitrogenase-mediated, we monitored the reaction under an argon (Ar) atmosphere as it is known that in the absence of N_2_ substrate, nitrogenase reduces protons to H_2_ (Hoffman et al., 2014). Indeed, the specific H_2_ photoproduction rate of the WT filaments under an Ar was about 7-fold higher compared to the N_2_ atmosphere (Fig. S9a). In the case of Δ*flv3A*, the yield of H_2_ photoproduction was strongly enhanced under an Ar and the production rate increased by around 10 times compared to N_2_ (Fig. S9a). In addition, a drastically decreased transcript abundance of *hoxH* in both Δ*flv1A* and Δ*flv3A* mutants compared to the WT (Fig. S9b), implied a negligible contribution of Hox to H_2_ production in Δ*flv3A* mutant. Collectively, these results provide evidence that the enhanced H_2_ production in the Δ*flv3A* mutant is mediated by nitrogenase.

**Figure 4.**
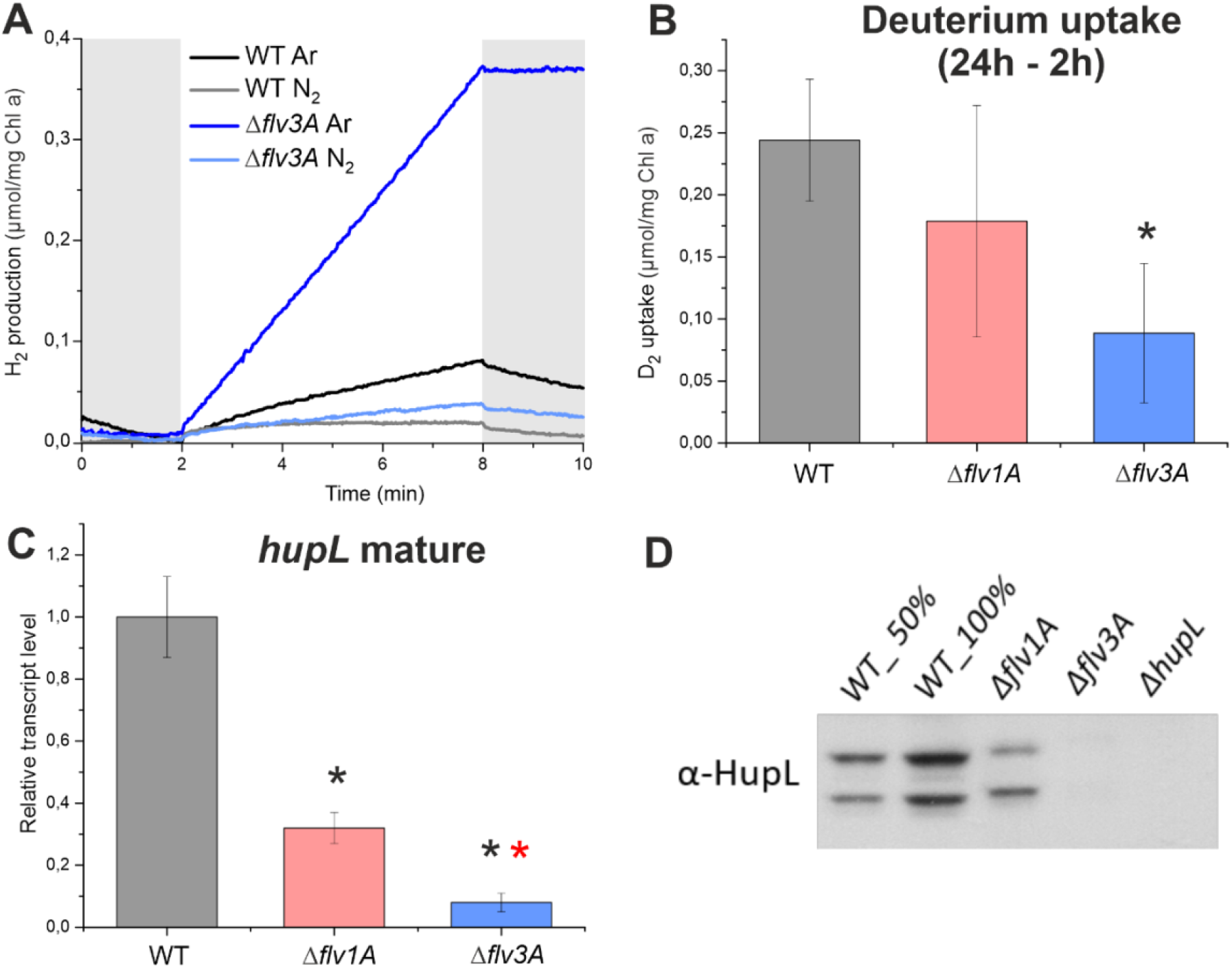
H_2_ metabolism of diazotrophic filaments of *Anabaena* WT, Δ*flvlA*, and Δ*flv3A*. (A)H_2_ production yield was monitored by a H_2_ electrode under an Ar or N_2_ atmosphere in the dark (grey areas) and under 800 μmol photons m^-2^ s^-1^ light. (B) Deuterium uptake by the filaments was calculated from the difference in D2 concentration between 2 h and 24 h after the injection in the vials initially flushed with Ar. (C) Relative transcript level of the mature *hupL*. (D) Immunodetection of HupL with a specific antibody. (E) Nitrogenase activity was measured using the acetylene reduction assay. Values are Mean ± SD, n = 3 biological replicates. Black asterisk indicates statistically significant differences compared to the WT (t-test, *P* < 0.05). Red asterisk indicates statistically significant differences compared to the Δ*flv1A* mutant (t-test, *P* < 0.05).

However, the observed decrease in nitrogenase activity (Table 1) did not correlate with an increase in H_2_ photoproduction in Δ*flv3A* (Fig. 3a). It is well known that the net nitrogenase-mediated production of H_2_ in heterocysts is strongly affected by the activity of uptake hydrogenase (Hup), which oxidizes H_2_ (Tamagnini et al., 2007). Therefore, it is conceivable that the impairment of the Hup function would account for the increased production of H_2_ in Δ*flv3A*. To examine the H_2_ fluxes, we traced the uptake of Deuterium (^2^H_2_, D_2_) by the WT, Δ*flv1A*, and Δ*flv3A*. Whilst WT and Δ*flv1A* filaments efficiently consumed D_2_, Δ*flv3A* showed a significantly lower capacity for D_2_ uptake (Fig. 4B). These observations confirmed that the impaired capacity of the Δ*flv3A* mutant to recycle H_2_ could be the reason for the increased accumulation of H_2_ observed in Δ*flv3A* mutant.

To better understand the molecular mechanism behind the defective H_2_ uptake of Δ*flv3A*, we analyzed the transcript and protein abundances of the large subunit of Hup (HupL). We found significant downregulation of the mature form of *hupL* transcript in both Δ*flv1A* and Δ*flv3A* mutants, in comparison to the WT (Fig. 4c). Importantly, *hupL* transcript level in Δ*flv3A* was significantly lower than in the *Δflv1A* mutant. Immunoblotting with specific antibody further revealed the lack of detectable HupL protein in Δ*flv3A*, a result comparable to the *hupL*-disrupted mutant (Δ*hupL*), while the Δ*flv1A* mutant showed only a lowered level of HupL relative to the WT (Fig. 4d). Taken together, these results demonstrate that the lack of HupL protein in heterocyst cells of *Δflv3A* is the reason behind the enhanced nitrogenase based H_2_ photoproduction observed for this mutant.

## Discussion

The heterocyst-forming cyanobacteria are considered one of the earliest forms of multicellular filaments in the history of life. Despite the extensive characterization of heterocyst differentiation, little is known about the co-regulation and interdependence of the two contrasting processes of N_2_ fixation and oxygenic photosynthesis occurring in heterocysts and vegetative cells, respectively. Under challenging environmental conditions, diazotrophic cyanobacteria must find an optimal balance between photochemical reactions and downstream processes that consume electrons in both cell types. In this work, we employed Δ*flv1A* and Δ*flv3A* mutants of *Anabaena* to examine the physiological significance of the vegetative cell-specific AnaFlv1A and AnaFlv3A proteins on the bioenergetic processes of diazotrophic cyanobacteria. Our results provide evidence that, in contrast to the *Synechocystis* homolog, AnaFlv3A can mediate moderate O_2_ photoreduction independently of AnaFlv1A and in coordination with AnaFlv2 and AnaFlv4 under LC conditions. Moreover, the vegetative-cell specific AnaFlv3A protein exhibits important link to the H_2_ metabolism inside the heterocyst, since the inactivation of this protein results in high H_2_ photoproduction even under ambient air. Nevertheless, we have demonstrated that both AnaFlv1A and AnaFlv3A proteins, presumably as hetero-oligomers, are required for efficient induction of the Mehler-like reaction during dark-to-light transitions, are crucial for photoprotection when light intensity rapidly fluctuates, and are likely needed for the activation of CO_2_ assimilation.

### In the absence of AnaFlv1A, AnaFlv3A can team up with AnaFlv2 and/or AnaFlv4 to mediate O_2_ photoreduction under LC conditions

In line with previous transcriptional analysis showing a decrease in the expression of both *flv1A* and *flv3A* in *Anabaena* WT upon the shift to diazotrophic conditions (Ermakova et al. 2014), the single deletions of AnaFlv1A or AnaFlv3A did not affect the diazotrophic growth of mutants under continuous illumination compared to the WT (Table 1). However, both AnaFlv1A and AnaFlv3A proteins are indispensable during sudden changes in light intensity, similar to their homologous proteins in other species (Fig. S1a, Allahverdiyeva et al., 2013; Gerotto et al., 2016; Jokel et al., 2018). Here, we have demonstrated that when both AnaFlv1A and AnaFlv3A proteins are expressed in WT filaments, the rate of the Mehler-like reaction is rapidly increased during the dark-to-light transition likely due to the activity of the AnaFlv1A/Flv3A hetero-oligomer (Fig. 3). Accordingly, the absence of either AnaFlv1A or AnaFlv3A delays rise in O_2_ photoreduction (Fig. 3a) resulting in over-reduction of the PQ pool upon illumination (Fig. 1a), causing a decrease in PSII yield (Fig. 1C) and impairment of PSI and Fd oxidation (Fig. 2). This phenotype is aggravated in the mutant lacking AnaFlv3A, which showed a stronger state 2-to-state 1 transition and more severe inability to oxidize PSI than the mutant lacking AnaFlv1A (Fig. 1 and Fig. 2). Differently from the *Synechocystis* Δ*flv1* mutant (Fig. S7), AnaFlv3A can promote O_2_ photoreduction in the *Anabaena* Δ*flv1A* mutant (Fig. 3a), resulting in only about 45% inhibition of steady-state O_2_ photoreduction and 35% decrease in Y(II) in *Anabaena* under LC growth conditions (Table S1). The near elimination of the steady-state O_2_ photoreduction in the Δ*flv1A/flv3A* double mutant under LC and the single mutants under HC conditions (where AnaFlv2 and AnaFlv4 are strongly downregulated) supports (i) functional AnaFlv3A/Flv2-4 oligomerization, and/or (ii) cooperation between the AnaFlv3A/Flv3A homo-oligomer and AnaFlv2/Flv4 hetero-oligomers. Accordingly, the strong impairment of O_2_ photoreduction in Δ*flv3A* might be due to the inability of AnaFlv1A to function as a homo-oligomer and/or cooperate with AnaFlv2/Flv4. It is worth emphasizing that both Δ*flv1A* and Δ*flv3A* mutants showed similarly enhanced accumulation of *flv2* and *flv4* transcripts (Fig. S4a). While the Δ*flv1A* mutant displayed WT-like *flv3A* transcript and protein levels, the Δ*flv3A* mutant showed an elevated *flv1A* transcript level compared to the WT (Fig. S4c). This shows that the inhibition of O_2_ photoreduction in Δ*flv3A* is not due to the downregulation of other FDPs. No contribution of the SynFlv3/Flv3 homo-oligomer in the Mehler-like reaction was observed *in vivo* (Mustila et al., 2016), contrary to previous *in vitro* studies suggesting a function of SynFlv3/Flv3 homo-oligomers in NAD(P)H-dependent O_2_ reduction (Vicente et al., 2002, Brown et al., 2019). Instead, a possible photoprotective function of SynFlv3/Flv3 homo-oligomers *via* an unknown electron transport network was proposed (Mustila et al., 2016). In *Anabaena* Δ*flv1A* mutant, AnaFlv3A/Flv3A homo-oligomers may, for example, be involved in controlling the cation homeostasis, which in turn may affect the reversible association of AnaFlv2/Flv4 hetero-oligomers with the thylakoid membrane, and consequently, their involvement in O_2_ photoreduction. It is also important to note that the oligomer formation scenario in the mutants might be different in *Anabaena* WT filaments. Overall, the obtained results suggest an important role for AnaFlv3A, but not AnaFlv1A, in mediating steady-state O_2_ photoreduction under diazotrophic LC conditions in an AnaFlv2/Flv4-dependent. Moreover, in LC but not in HC conditions, the lack of both AnaFlv1A and AnaFlv3A resulted in a more severe delay in induction of O_2_ evolution during dark-to-light transition in comparison to the lack of AnaFlv3A only (Fig. 3). This suggests that AnaFlv1A may also function in coordination with AnaFlv2/4 independently of Flv3A in an unknown role that facilitates photosynthetic electron transport. Understanding the exact functions of AnaFlv2 and/or AnaFlv4 in these processes and their interactions with AnaFlv1A and AnaFlv3A requires further investigation.

Even though AnaFlv1A and AnaFlv3A contribute to the Mehler-like reaction to different extents, both Δ*flv1A* and Δ*flv3A* mutants exhibited reduced CCM activity, as deduced from lowered initial peaks in CO_2_ uptake rate during dark-to-light transition and reduction of steady-state CO_2_ uptake in comparison to WT (Fig. S8b, Table S2). This is likely to result from impaired energization of CCM in the absence of AnaFlv1A or AnaFlv3A. SynFlv1 and SynFlv3 have been shown to have a crucial role in the generation of *pmf* during the dark-to-light transition (Nikkanen et al., 2020), comparable to that of FLVA/B in *P. patens* (Gerotto et al., 2016) and *C. reinhardtii* (Chaux et al., 2017). Moreover, the *pmf* generated by FDPs and CET has been recently shown to be important for inducing and maintaining CCM activity in *C. reinhardtii* (Burlacot et al., 2021). We hypothesize that AnaFlv1A/Flv3A hetero-oligomer is required to rapidly induce the Mehler-like reaction, likely being important for the generation of *pmf* and possibly for induction of CCM activity during the dark-to-light transition. The molecular mechanism of the FDP-dependency of the CCM requires further investigation however, as the mechanisms of CCM differ between *Chlamydomonas* and cyanobacteria (Price et al 2008). Moreover, in *Synechocystis* mutants lacking Flv1/3 *pmf* generation during the first minute of dark-to-light transitions is severely impaired, CCM induction is largely unaffected at least in standard conditions (Nikkanen et al., 2020). In *Anabaena* however, both Δ*flv1A* and Δ*flv3A* strains demonstrated impaired induction of CCM (Fig. S8b), suggesting that *Anabaena* may differ from *Synechocystis* in the extent to which CCM induction is *pmf*-dependent.

Compelling evidence has recently been provided for dynamic coordination and functional redundancy between NDH-1 and SynFlv1/Flv3, jointly contributing to efficient oxidation of PSI in *Synechocystis* (Nikkanen et al., 2020) and in *Physcomitrella patens* (Storti et al., 2020a, 2020b). NDH-1-mediated cyclic electron transport (CET) in *Anabaena* could also partially compensate for a lack of AnaFlv1A and AnaFlv3A as evidenced by a stronger F_0_ rise observed in both mutants (Fig. S3a). Unlike *Synechocystis* cells, *Anabaena* filaments express orthologs of plastid terminal oxidase (PTOX, *all2096*) (McDonald et al., 2003). It has been proposed that in *C. reinhardtii* and vascular plants, PTOX functions as an electron valve directing electrons from plastoquinol to O_2_, thereby controlling the redox state of the PQ pool (Stepien and Johnson, 2018, Saroussi et al., 2019; Nawrocki et al., 2019) and being involved in diverse metabolic processes such as the regulation of CET, state transition and carotenoid biosynthesis (Nawrocki et al., 2019). We cannot exclude possible contribution of PTOX to the residual O_2_ photoreduction observed in the Δ*flv3A* mutant (Fig. 3A), and/or as a sensor of the redox state of the PQ pool and regulator of NDH-dependent CET, thus limiting electron pressure on the acceptor-side of PSI (Bolte et al., 2020).

### Inactivation of AnaFlv3A leads to enhanced nitrogenase-based H_2_ photoproduction under oxic conditions

Demonstration of elevated photoproduction of H_2_ gas in diazotrophic filaments lacking vegetative cell-specific AnaFlv3A under oxic and microoxic conditions (Fig. 4) provided intriguing information about bioenergetic interdependence between vegetative cells and heterocysts. The heterocyst-originated production of H_2_ in the Δ*flv3A* mutant was rapidly induced upon exposing the filaments to light and occurred concomitantly with the evolution of O_2_ in neighbouring vegetative cells (Fig. 3). Moreover, the rate of H_2_ photoproduction in the Δ*flv3A* mutant responded positively to an increase in CO_2_ availability (Fig. 3b).

In the absence of N_2_, the main substrate for nitrogenase, all electrons can be directed to H_2_ production (Hoffman et al., 2014) allowing a less costly reaction, whereby only 4 moles of ATP are required to produce one mole of H_2_. In this work, the removal of N_2_ substrate (by replacement with Ar) led to a 10-fold increase of H_2_ photoproduction rate in Δ*flv3A*, demonstrating the occurrence of nitrogenase-dependent H_2_ photoproduction in this mutant (Fig. 4a). A recent report suggested that overexpressing Flv3B lead to more stable microoxic conditions inside the heterocysts, notably increasing the H_2_ production yield, presumably *via* the bidirectional hydrogenase Hox (Roumezi et al., 2020). In contrast to the unidirectional production of H_2_ by nitrogenase, Hox catalyzes the reversible reduction of protons to H_2_ (Bothe et al., 2010). We do not consider the contribution of Hox to the photoproduction of H_2_ by the Δ*flv3A* mutant, as the net production does not fit with the bidirectional nature of the enzyme. Moreover, significant downregulation in the Δ*flv3A* mutant of transcripts from *hoxH*, encoding one of the subunits (Fig. S9b) further supports this assumption. Altogether, these results indicate that the increased light-induced H_2_ photoreduction of the Δ*flv3A* mutant is mediated by nitrogenase activity.

Strikingly, it turned out that the increase in H_2_ photoproduction yield in the Δ*flv3A* mutant was due to significant downregulation of HupL, the large subunit of the uptake hydrogenase, evidenced both at the transcript and protein levels (Fig. 4c, 4d). The absence of functional Hup suppressed the H_2_ recycling pathway (Fig. 4b) and caused the release of H_2_, photoproduced by nitrogenase, from the heterocysts of Δ*flv3A* filaments (Fig. 3, Fig. 4a). Thereby, our results highlight a regulatory network between the two metabolic processes in different compartments: The Flv3A-mediated metabolic processes in the vegetative cells and the H_2_ metabolism in the heterocysts. It is likely that the redox state of the PQ pool in vegetative cells, affected by the activity of Flv3A, has a regulatory role on the H_2_ metabolism in heterocysts. However, the nature of the molecular signal from reduced PQ that ultimately regulates gene expression in heterocysts remains unknown. The redox state of the PQ pool in likely correlates with the availability of soluble reducing cofactors in the cytosol of vegetative cells, and while evidence in *Anabaena* is lacking, those cofactors may be interchanged between vegetative cells and heterocysts, inducing changes in metabolism and gene expression. A majority of the NADPH needed for the nitrogen metabolism in heterocysts is understood to be derived from the oxidative pentose phosphate pathway breaking down carbohydrates imported from vegetative cells, (Cumino et al., 2007) but it is plausible that more direct exchange of cofactors also occurs, analogously to the malate redox shuttle between cytosol and the chloroplast in plants and algae. Nevertheless, the molecular mechanism underlying this regulatory network between different cell types needs further elucidation.

Taken together, our results demonstrate that similarly to SynFlv1 and SynFlv3, both vegetative-cells specific AnaFlv1A and AnaFlv3A are indispensable under harsh FL conditions regardless of nitrogen or CO_2_ availability, most likely maintaining sufficient oxidation of the photosynthetic electron transport chain by catalysing the Mehler-like reaction as AnaFlv1A/Flv3A hetero-oligomers. Under LC, AnaFlv3A is able to perform moderate O_2_ photoreduction in coordination with AnaFlv2 and AnaFlv4 proteins and independently of AnaFlv1A. AnaFlv3A may either stimulate the activity of AnaFlv2/Flv4 hetero-oligomers indirectly via an unknown function, or participate in forming functional hetero-oligomers with AnaFlv2 and Flv4. The deletion of AnaFlv3A was concomitant with the downregulation of the heterocyst-specific Hup enzyme resulting in increased bioproduction of H_2_. This novel regulatory network between the photosynthesis and diazotrophic metabolism might represent an unexploited source for the future of biotechnological applications.

## Acknowledgements

We thank Prof. Paula Tamagnini for the HupL antibody.

## Authors contribution

Y.A. conceived the study. A.S-S., L.N., G.T., M.E., S.K., and Y.A. designed the research. A.S-S. performed most of the experiments. M.E. performed growth characterization of the mutants. L.N. performed KLAS-NIR, S.K measured H_2_ production using the electrode. M.H. performed Deuterium uptake experiment. G.T performed RT-qPCR experiments and J.W constructed an independent Δ*flv3A* mutant. EMA provided resources. A.S.S. drafted the manuscript and all authors revised and approved it.

## Data availability

The data that support the findings of this study are available from the corresponding author upon reasonable request.

