## Supplementary material for "Flv3A facilitates O_2_ photoreduction and affects H_2_ photoproduction independently of Flv1A in diazotrophic *Anabaena* filaments": Suppl Material

**Supplemental materials**

### Material and methods

##### Determination of P700 and Fd redox changes from near-infrared absorbance

Redox change kinetics of P700 and Fd were deconvoluted from the four difference signals using differential model plots (model spectra) (Supplemental Figure 10) that were measured for *Anabaena* using protocols described earlier (Theune et al., 2021) with the modification for the Fd model spectrum, we used the Δ*flv1A*/Δ*flv3A* mutant instead of anoxic conditions to impair the Mehler-like reaction (see Figure 2) and to sufficiently slow down the re-oxidation Fd. For P700 and plastocyanin (PC) model spectra, we used WT *Anabaena* filaments. Due to low signal quality, the PC traces were omitted from Figure 2. As noted for *Synechocystis* earlier (Theune et al., 2021), it is likely that the redox kinetics of P700 and PC in *Anabaena* may be closely related, thus making it difficult to extract a PC signal of large magnitude.

### Figures

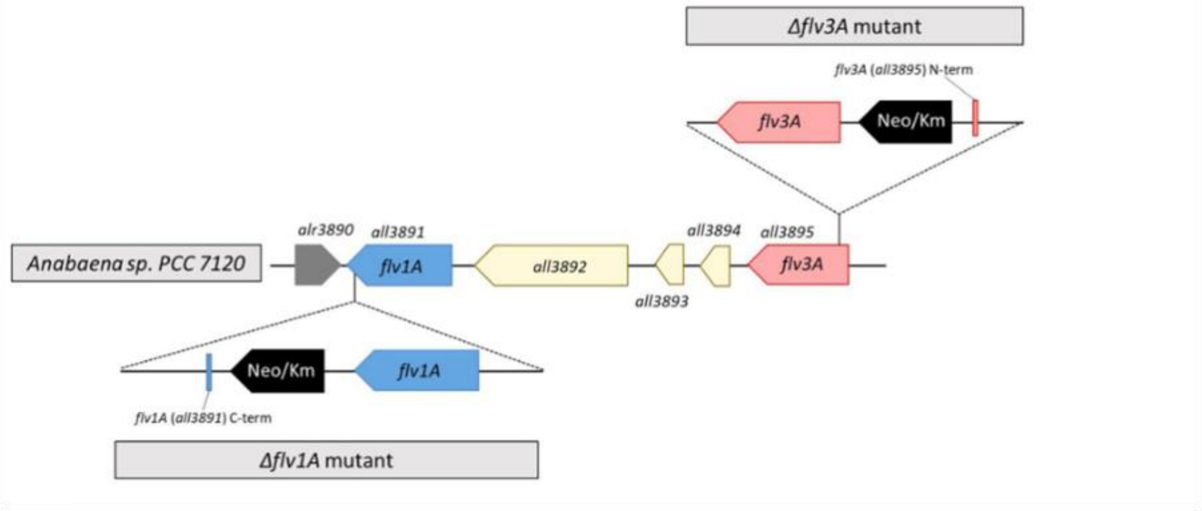


**Supplemental Figure 1.** The genomic structure of *Anabaena* Δ*flv1A* and Δ*flv3A* mutants used in this work.

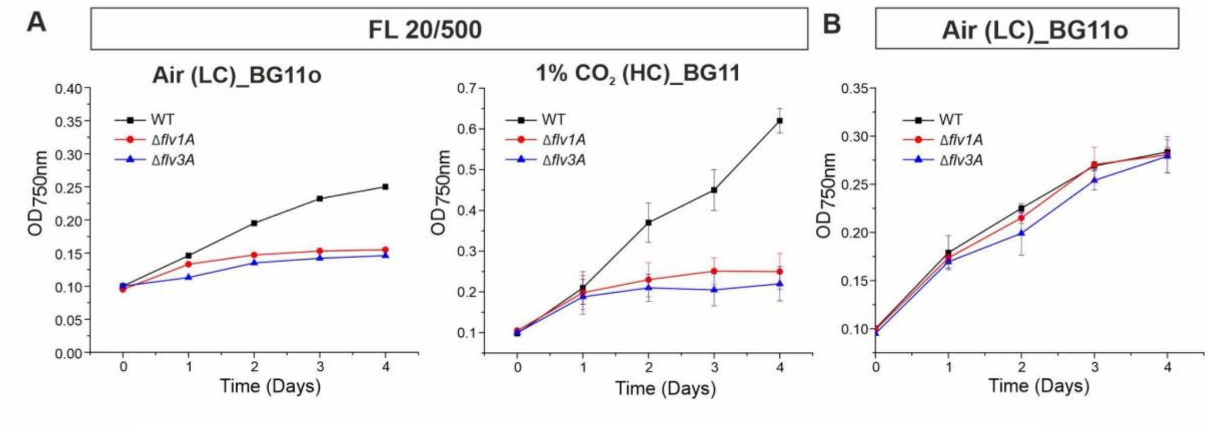


**Supplemental Figure 2**. Growth characterization of *Anabaena* WT, Δf*lv1A* and Δ*flv3A* filaments. (a) Growth curves of filaments grown under severe fluctuating light (FL 20/500) through bubbling with air or 1% $CO_2$. (b) Diazotrophic growth in $BG11_0$ and air bubbling.

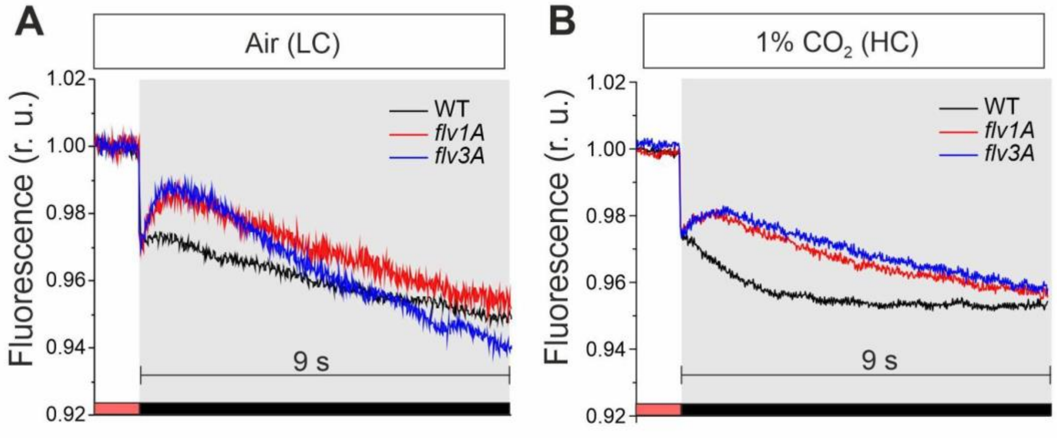


**Supplemental Figure 3. F0 rise of *Anabaena* WT, Δ*flv1A* and Δ*flv3A*.** Transient F0 rise was monitored after switching off the red actinic light (50 µmol photons $m^{-2}$ $s^{-1}$, red bars) for filaments grown under (A) air or (B) 1% $CO_2$ ($F_0$ rise was normalised to $F_s$ to facilitate comparison of the kinetics).

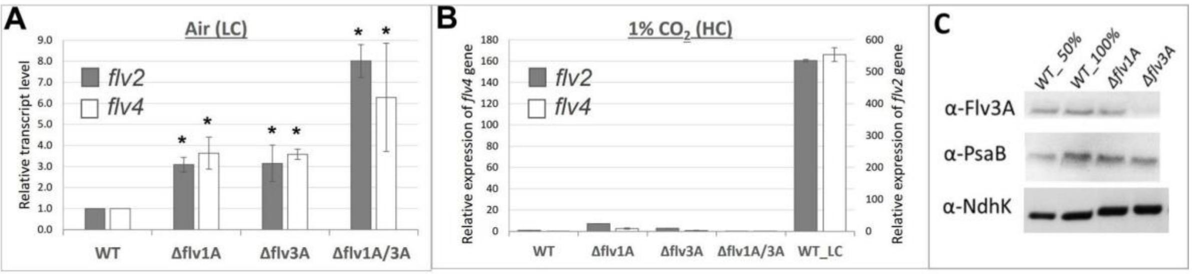


**Supplemental Figure 4. Analyses of transcript and protein abundance in the diazotrophic WT, Δ*flv1A* and Δ*flv3A* filaments.** Relative transcript abundance of *flv2* and *flv4* grown under (A) LC (air) and (B) HC (1% $CO_2$). Values are Mean ± SD, n = 3 biological replicates. Asterisks indicate statistically significant differences between the mutants and WT (t-test, P < 0.05). (C) Immunodetection of Flv3A, PsaB and NdhK from diazotrophic filaments grown under LC.

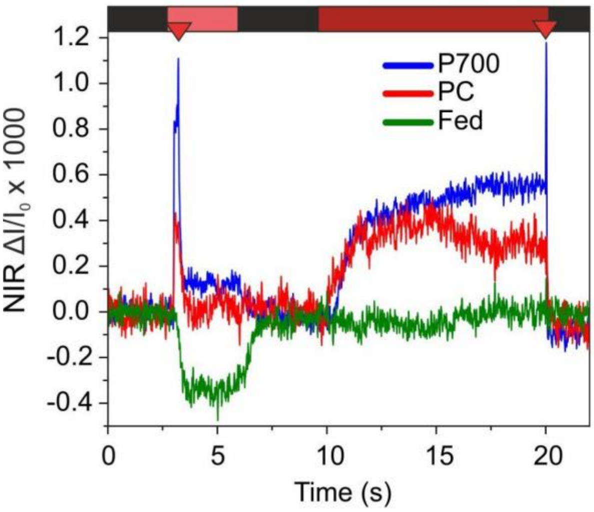


**Supplemental Figure 5. DUAL-KLAS-NIR kinetics of P700, PC and Fd in *Synechocystis* Δ*flv1* mutant.**
The Δ*flv1* mutant was grown in ambient $CO_2$ level and illuminated with 50 µmol photons $m^{-2}s^{-1}$ for 4 days in BG-11 pH 7.5, after which cells were harvested and Chl a concentration adjusted to 10 µg $ml^{-1}$ with fresh BG-11. Maximum amplitudes of PC, P700 and Fd in Δ*flv1* cells were determined with the NIRMAX script of the DUAL-KLAS-NIR software. 3 s illumination of red AL (200 µmol photons $m^{-2}s^{-1}$), with a saturating pulse (depicted by a red triangle) after 200 ms of illumination to fully reduce the Fd pool. After 4 s of darkness thereafter, cells were illuminated with far red light for 10 s with a saturating pulse in the end to fully oxidize P700.

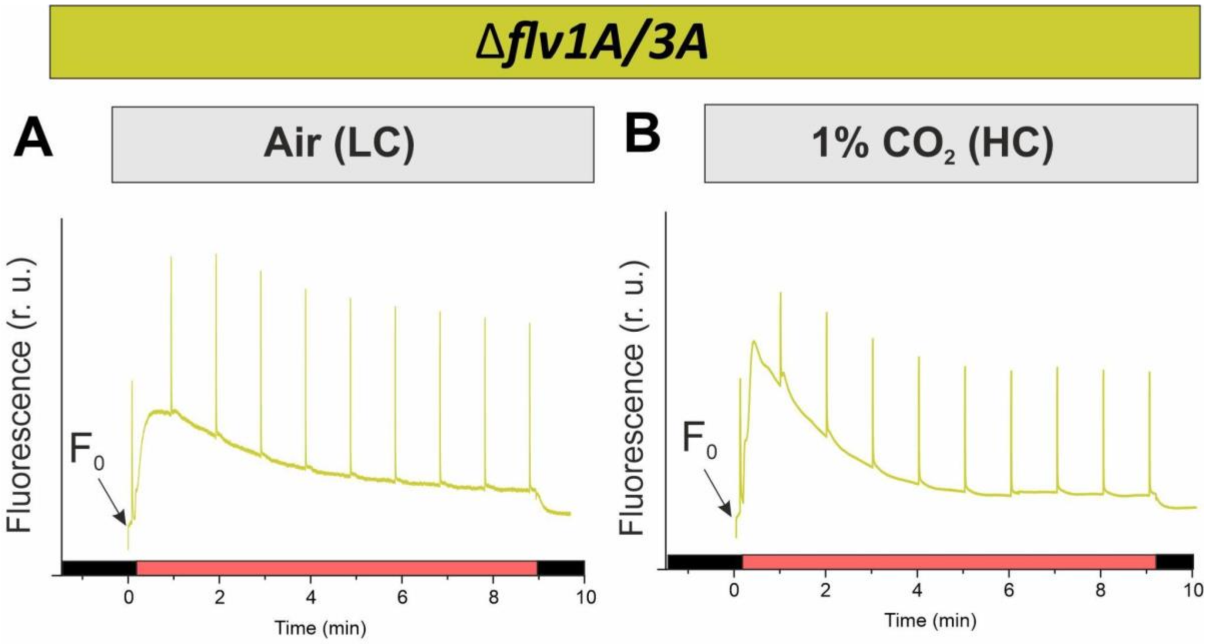


**Supplemental Figure 6. Fluorescence induction curves of diazotrophic *Anabaena* Δ*flv1A/flv3A* cultivated under LC or HC.** (A, B) The kinetics are representative of three biological replicates. Cells were dark acclimated for 10 min before the measurement.

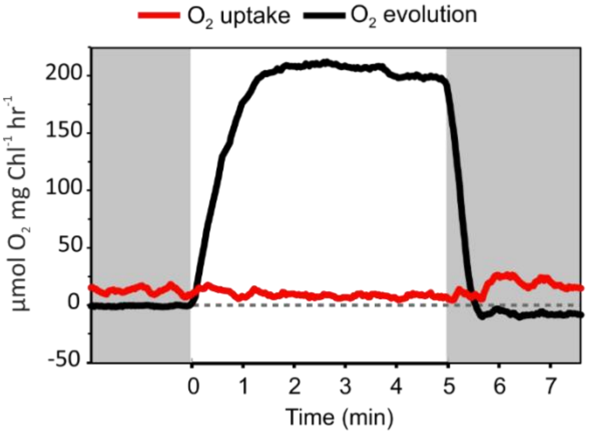


**Supplemental Figure 7. $O_2$ exchange rates of the *Synechocystis* Δ*flv1* mutant.** Experimental cultures were grown in BG11 medium in air level of $CO_2$ and illuminated with 50 μmol photons $m^{-2}$ $sec^{-1}$ light for 4 days. After this the cells were harvested and chlorophyll (Chl) a concentration adjusted to 10 μg $ml^{-1}$ with fresh BG-11. Cells were dark-adapted for 15 min, and gas exchange was monitored by MIMS over a 5-min llumination period with 500 μmol photons $m^{-1}$ $sec^{-2}$ of white actinic light. Before the measurements, samples were supplemented with 1.5 mM $NaHCO_3$.

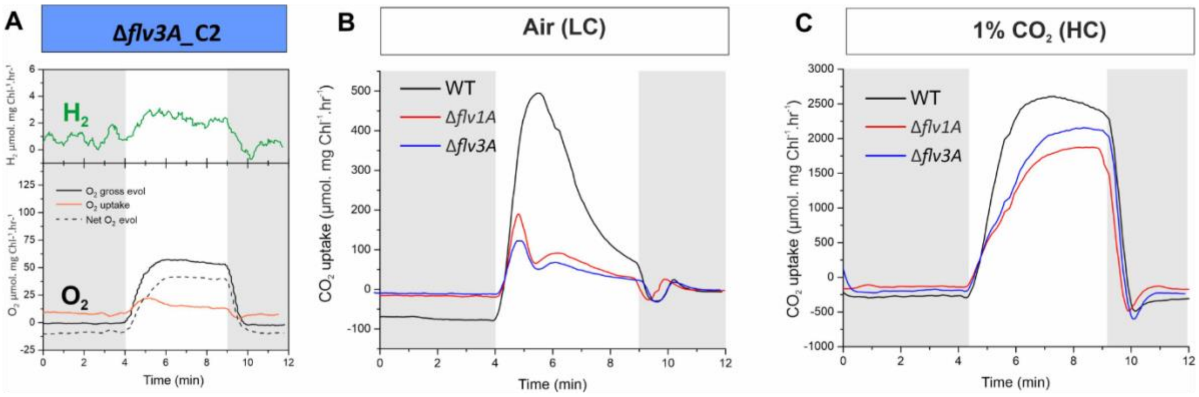


**Supplemental Figure 8. Gas exchange analysis of the diazotrophic *Anabaena* filaments.** (A) $O_2$ and $H_2$ exchange rates of a second independent mutant colony Δ*flv3A*_C2 grown under LC. (B, C) $CO_2$ uptake rates of WT, Δ*flv1A*, and Δ*flv3A* filaments grown under LC (B) or HC (C). Changes of the $CO_2$ concentration were recorded simultaneously to the measurements in Fig 2. Gas exchange rates are presented as μmol mg Chl $a^{-1}$ $h^{-1}$. Mean ± SD, n = 2-3 biological replicates.

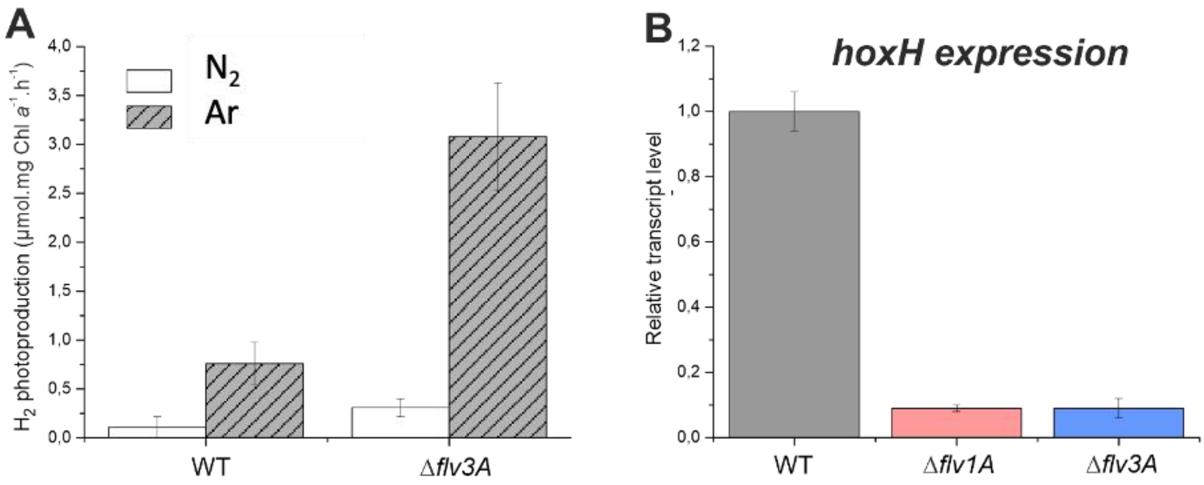


**Supplemental Figure 9. $H_2$ metabolism in diazotrophic filaments of *Anabaena* WT and Δ*flv1A* and Δ*flv3A* mutants** (A) Specific $H_2$ photoproduction rates under an Ar or $N_2$ atmosphere. (B) Transcript abundance of *hoxH*.

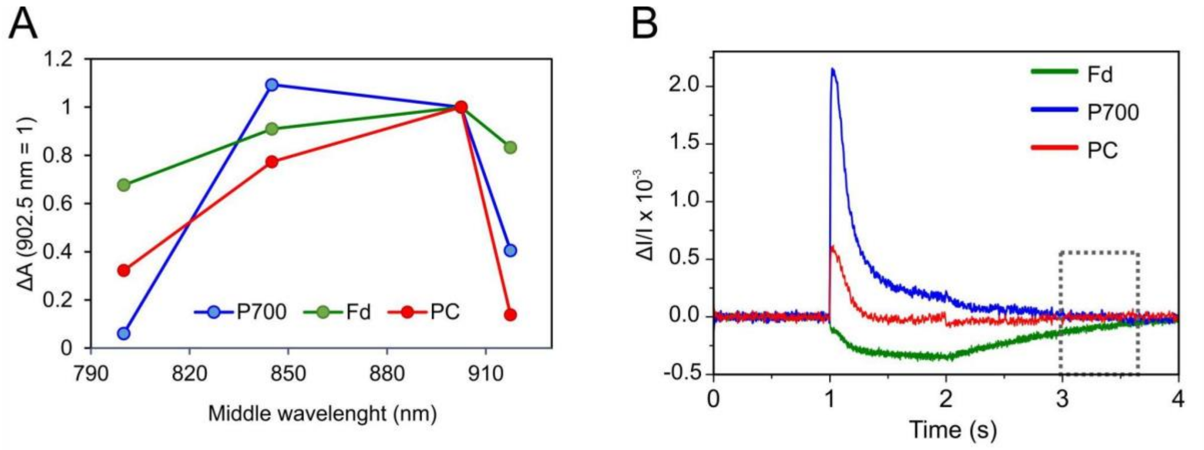


**Supplemental Figure 10. Differential model blots (DMPs) for deconvolution of PC,P700, and Fd signals with the DUAL-KLAS-NIR spectrometer.** (A) DMPs determined from WT (P700 and PC) and Δflv1A/Δflv3A (Fd) Anabaena cells according to protocols described by Theune et al. (2021), with the exception that the Δflv1A/Δflv3A mutant was used instead of anoxic conditions to inhibit the Mehler-like reaction and thus delay reoxidation of Fd sufficiently to determine the Fd DMP. The values are normalized to the ΔA value at the middle wavelength at 902.5 nm. (B) Deconvoluted traces from the determination of the Fd DMP. Δflv1A/Δflv3A cells were illuminated for 1 s at 1350 µmol photons $m^{-2}$ $s^{-1}$ to fully reduce the Fd pool. The illumination was repeated 10 times with a 30 s interval in between measurements, and averaged traces from the 10 measurements are shown. The dashed rectangle indicates the time window where only Fd redox changes occur and which was used to determine the Fd DMP.

**Supplemental Table 1.** Oligonucleotide sequences used for qPCR.

| **Gene name** | **Forward primer (5′ →3′)** | **Reverse primer (3′ →5′)** |
|---|---|---|
| *flv2* (*all4444*) | cgacttttgcccaaacttta | gatcgccatcataattcctg |
| *flv4* (*all4446*) | ctgctattcgctgtttggat | ttcactaagccgctatggtc |
| *hupL_mature* | agtagccgcttctacgatga | acccaaccacacaggttcta |
| *hoxH* | gggacaaatcctccaatccc | tttgctcctcccaacacttc |
| *rnpB* | ggactaggggttggggact | acgagggcgattatctatctg |

**Supplemental Table 2.** The rates of $CO_2$ and $O_2$ exchange in WT, Δ*flv1A*, and Δ*flv3A* filaments grown under air (LC) or in the air supplemented with 1% $CO_2$ (HC). Gas exchange rates are presented as µmol gas mg Chl $a^{-1}$ $h^{-1}$. Mean ± SD, n = 2-3 biological replicates, N.D., not detected. Asterisks indicate statistically significant differences compared to the WT (t-test, $P < 0.05$).

| | Dark respiration | Gross $O_2$ evolution (steady-state) | Light-induced $O_2$ uptake (max. peak / steady-state) | Net $O_2$ evolution | $CO_2$ uptake (max. peak / steady- state) | Ratio Gross $O_2$ evolution/ $O_2$ uptake (steady-state) |
|---|---|---|---|---|---|---|
| **Air (LC)** | | | | | | |
| **WT** | 10.6±2.7 | 104.4±2.9 | 34.6±0.2/ 28.4±1.4 | 70±0.7 | 485.3±48.1/ 65.0±19.6 | 3.7±0.3 |
| **Δ*flv1A*** | 8.5±1.5 | 101.1±8.3 | 22.2±2.8/ 15.5±1.2* | 77.1±8.6 | 190.0±32.6*/ 29.1±10.9* | 6.5±0.0 |
| **Δ*flv3A*** | 8.2±0.1 | 73.7±2.3 | 15.6±0.1/ 4.3±0.2* | 61.1±2.2 | 112.5±21.3*/ 20.6±9.2* | 17.0±0.2 |
| **1% $CO_2$ (HC)** | | | | | | |
| **WT** | 11.0±2.4 | 122.2±14.8 | 32.9±1.4/ 30.6±1.5 | 80.7±11.0 | 2523.8±300.7 | 4.0±0.3 |
| **Δ*flv1A*** | 8.8±1.5 | 65.9±7.3 | N.D.* | 56.5±9.3 | 1842.6±201.3* | -- |
| **Δ*flv3A*** | 8.4±2.1 | 76.82±6.45 | N.D.* | 69.2±4.0 | 2116.9± 206.8 | -- |
